# Adaptive arousal regulation: Pharmacologically shifting the peak of the Yerkes-Dodson curve by catecholaminergic enhancement of arousal

**DOI:** 10.1101/2024.09.11.612196

**Authors:** Lola Beerendonk, Jorge F. Mejías, Stijn A. Nuiten, Jan Willem de Gee, Jasper B. Zantvoord, Johannes J. Fahrenfort, Simon van Gaal

## Abstract

Performance typically peaks at moderate arousal levels, consistent with the Yerkes-Dodson law, as confirmed by recent human and mouse pupillometry studies. Arousal states are influenced by neuromodulators like catecholamines (noradrenaline; NA and dopamine; DA) and acetylcholine (ACh). To explore their causal roles in this law, we pharmacologically enhanced arousal while measuring human decision-making and spontaneous arousal fluctuations via pupil size. The catecholaminergic agent atomoxetine (ATX) increased overall arousal and shifted the entire arousal-performance curve, suggesting a relative arousal mechanism where performance adapts to arousal fluctuations within arousal states. In contrast, the cholinergic agent donepezil (DNP) did not affect arousal or the curve. We modeled these findings in a neurobiologically plausible computational framework, showing how catecholaminergic modulation alters a disinhibitory neural circuit that encodes sensory evidence for decision-making. This work suggests that performance adapts flexibly to arousal fluctuations, ensuring optimal performance in each and every arousal state.

## Introduction

Behavioral performance typically peaks during mid-level arousal – a relationship described by the classic Yerkes-Dodson law and recently confirmed in human and mouse pupillometry studies^1–6^. Arousal fluctuates continuously across several time scales (e.g., days, hours, minutes, seconds) and these fluctuations are primarily influenced by the catecholaminergic (CA) and cholinergic (ACh) neurotransmitter systems^5,7–9^, which regulate the balance between sympathetic (fight or flight mode) and parasympathetic nervous system activity (rest and digest mode). The neuromodulator systems that regulate arousal also modulate pupil dilation^5,7,10–15^, making non-luminance-mediated changes in pupil size a robust marker of spontaneous fluctuations of arousal, often referred to as “pupil-linked arousal”^5,16–18^.

Given that pupil size reflects the combined influence of various neuromodulatory systems (besides ACh and catecholamines e.g., also serotonin and orexin)^7,10,11,13,14,19^, previous work using pupil-linked arousal could not effectively distinguish the individual contributions of each neuromodulator to the Yerkes-Dodson law. Precise manipulation of these systems is crucial for unraveling their distinct roles in arousal-performance dynamics. Clarifying these mechanisms is not just a fundamental question in neuroscience, it also has significant implications for treating conditions like ADHD, anxiety, and sleep disturbances, which all involve arousal dysregulation^20,21^. Additionally, disentangling the influence of neuromodulators on performance could enhance human performance in high-stakes environments (e.g., aviation, emergency response) by informing strategies to optimize arousal levels.

Here we pharmacologically increase neuromodulation using two different drugs that each enhance one main neuromodulatory driver of cortical arousal^5,8,22^ to assess their relative roles in the inverted U-shaped arousal-performance relationship. The first drug, Donepezil (DNP), is a selective cholinesterase inhibitor that increases ACh levels by hindering the breakdown of acetylcholine^23^. The second drug, Atomoxetine (ATX), increases noradrenaline (NA) and dopamine (DA) levels by blocking their reuptake into the presynaptic terminal via NA transporters, serving as a relatively selective noradrenaline reuptake inhibitor (as both NA and DA are CAs, we refer to ATX as a catecholaminergic drug)^24^. Both neuromodulators have been shown to affect overall performance in simple decision-making tasks (ACh:^25–27^, catecholamines:^26,28–32)^. By comparing the relationship between pupil-linked arousal fluctuations and performance in these enhanced neuromodulation states to a placebo control condition (PLC), we aim to clarify the roles of catecholamines and ACh in the pupil-linked arousal-performance relationship.

In addition, this pharmacological approach allows us to address another key open question about the Yerkes-Dodson law: whether it is absolute (and fixed) or relative (and adaptive). The common conception is that there is a fixed relationship between arousal and performance, with optimal performance during moderate arousal states, and inferior performance during states of low and high arousal. In this scenario, performance depends on the absolute arousal level and therefore the Yerkes-Dodson curve spans the entire arousal spectrum (absolute arousal scenario). Consequently, once arousal surpasses the moderate (“optimal”) level, performance will keep declining with increasing arousal, due to the selective sampling of only the right (descending) half of the curve (**Figure 1A**). Alternatively, when an organism moves from a relaxed context to a more stressful context, the neural circuitry may adapt to these new circumstances. In this relative arousal scenario, the relationship between arousal fluctuations and task performance is inverted U-shaped within each context (**Figure 1B)**. In other words, even though the position of the pupil-linked arousal-performance relationship might change due to overall arousal changes, its shape remains intact. Such flexible adjustments to environmental conditions, reminiscent of (divisive) normalization^33–35^, have been observed in other domains in neuroscience as well. For example, neuronal responses in V1 to a stimulus can adapt based on the average contrast in the visual field^34^, and in mouse hippocampal network activity, neurons can dynamically scale their firing range depending on the size of the surrounding environment of the animal^36^. For the first time in humans, we combine pharmacologically induced arousal state manipulations with simultaneous and continuous measurement of pupil-linked arousal fluctuations and behavioral psychophysics to arbitrate between the absolute and relative arousal scenario. In addition, we use computational modelling to gain a better understanding of the empirical results.

**Figure 1.**
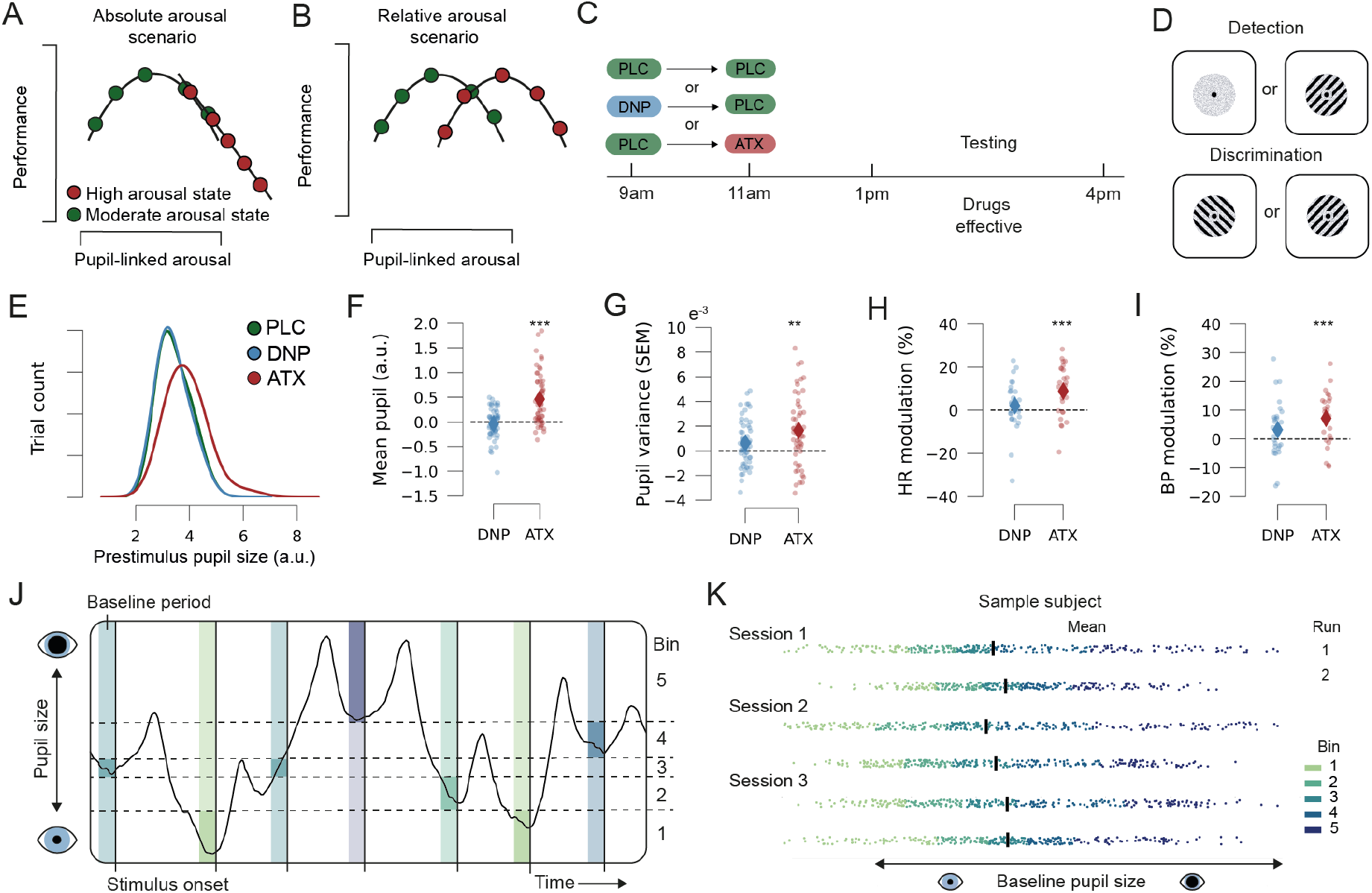
Scenarios, tasks, arousal modulation, and pupil analysis. (A) Schematic of the absolute and (B) relative arousal scenario. (C) Outline of a session including drug administration scheme. (D) Schematic stimuli of the detection and discrimination tasks. (E) Histogram of prestimulus pupil size split by drugs. (F) Mean prestimulus pupil size modulation for DNP and ATX as compared to PLC. (G) Prestimulus pupil variance (standard error of the mean: SEM) modulation for DNP and ATX as compared to PLC. (H) Heart rate (HR) modulation and (I) Blood pressure (BP) modulation for DNP minus PLC and ATX minus PLC (referred to as ATX and DNP respectively on the x-axis). (J) Example of the continuous pupil signal of seven trials with baseline windows [-.5s to 0s to stimulus onset] divided over five bins. Colors indicate pupil size bins, see legend in K. (K) Example data of three sessions of the discrimination task of a sample participant. Bins are determined for each task run (block) separately. Each dot represents the baseline pupil size of a trial, and the colors indicate bin membership (1-5). Black vertical lines indicate mean pupil size of each run. ***p<.001, **p<.01, statistics in text.

## Results

Participants (N=28) took part in three all-day (9am-4pm) experimental sessions in which they either received PLC, DNP (5mg), or ATX (40mg) in randomized order (**Figure 1C**). The administration scheme ensured that blood concentration levels for DNP and ATX peaked when the tasks commenced (see Methods). All participants performed four different visual decision-making tasks: two Gabor detection tasks (presence/absence judgement) and two Gabor orientation discrimination tasks (left/right tilt judgement, **Figure 1D**). We recorded on average 6337 trials per participant, resulting in a total of 177460 trials. Performance was matched across all tasks and titrated to 75% correct (mean accuracy across tasks was 75.7% for the PLC condition).

First, we assessed whether the pharmacological agents ATX and DNP had increased physiological arousal as compared to PLC. To index arousal, we measured prestimulus pupil size, prestimulus pupil size variation, heart rate (HR) and mean arterial blood pressure (BP). We observed a clear increase in mean prestimulus pupil size, indicative of a heightened arousal state, after ingestion of ATX as compared to PLC (t(27)=4.78, p<.001; **Figure 1E-1F**; more statistical tests in the **Supplements**). In addition, ATX increased the range (i.e., variance) of arousal fluctuations, evidenced by a larger variation in prestimulus pupil size (t(27)=3.43, p=.002; **Figure 1G**). Lastly, ATX increased heart rate and blood pressure (HR: t(27)=4.11, p<.001; BP: t(27)=4.33, p<.001; **Figure 1H-I**). DNP did not affect any of these measures (more details in the **Supplements**, also see ^31^). Previous non-clinical studies that report physiological responses of 5mg DNP^37,38^ also did not observe consistent subjective or physiological effects of DNP. However, even in the absence of such autonomic markers, 5mg DNP has been shown to induce changes in task behavior and associated neural activity patterns^25,27,31,37–40^. Therefore, we still performed the intended analyses for DNP.

Next, we confirmed that the arousal-performance relationship was indeed inverted U-shaped in the PLC condition, as we previously showed for both visual as well as auditory sensory input^1^. Identical to our previous study, we used mean pupil size in the 500ms leading up to target onset as a proxy for arousal. We initially assigned each trial to one of twenty equally populated bins based on prestimulus pupil size (**Figure 1J**; visualized for five bins). We performed this binning procedure for each run of all tasks separately, thereby focusing on arousal fluctuations that occur over the course of single experimental runs (i.e., blocks, **Figure 1K**). For each pupil bin we calculated Signal Detection Theoretic sensitivity (d’^41^) and mean reaction times (RT). Note that an increase in performance is equivalent to a decrease in RT, which means that the expected relationship between arousal and RT is U-shaped (not inverted)^1^. We used linear mixed models and probabilistic model comparison to assess whether the pupil-performance relationship was linear or quadratic. A difference in AIC^42^ or BIC^43^ values of more than 10 (i.e., ΔAIC or ΔBIC>10) is considered evidence that the winning model captures the data significantly better^44^. Both ΔAIC and ΔBIC were strongly in favor of a quadratic pupil-performance relationship for d’ and RT (all ΔAIC>20.1, all ΔBIC>15.8; **Figure 2A and D**, green dots; **Supplementary table 2**) in the PLC condition, in line with previous work^1,16–18,45,46^.

**Figure 2.**
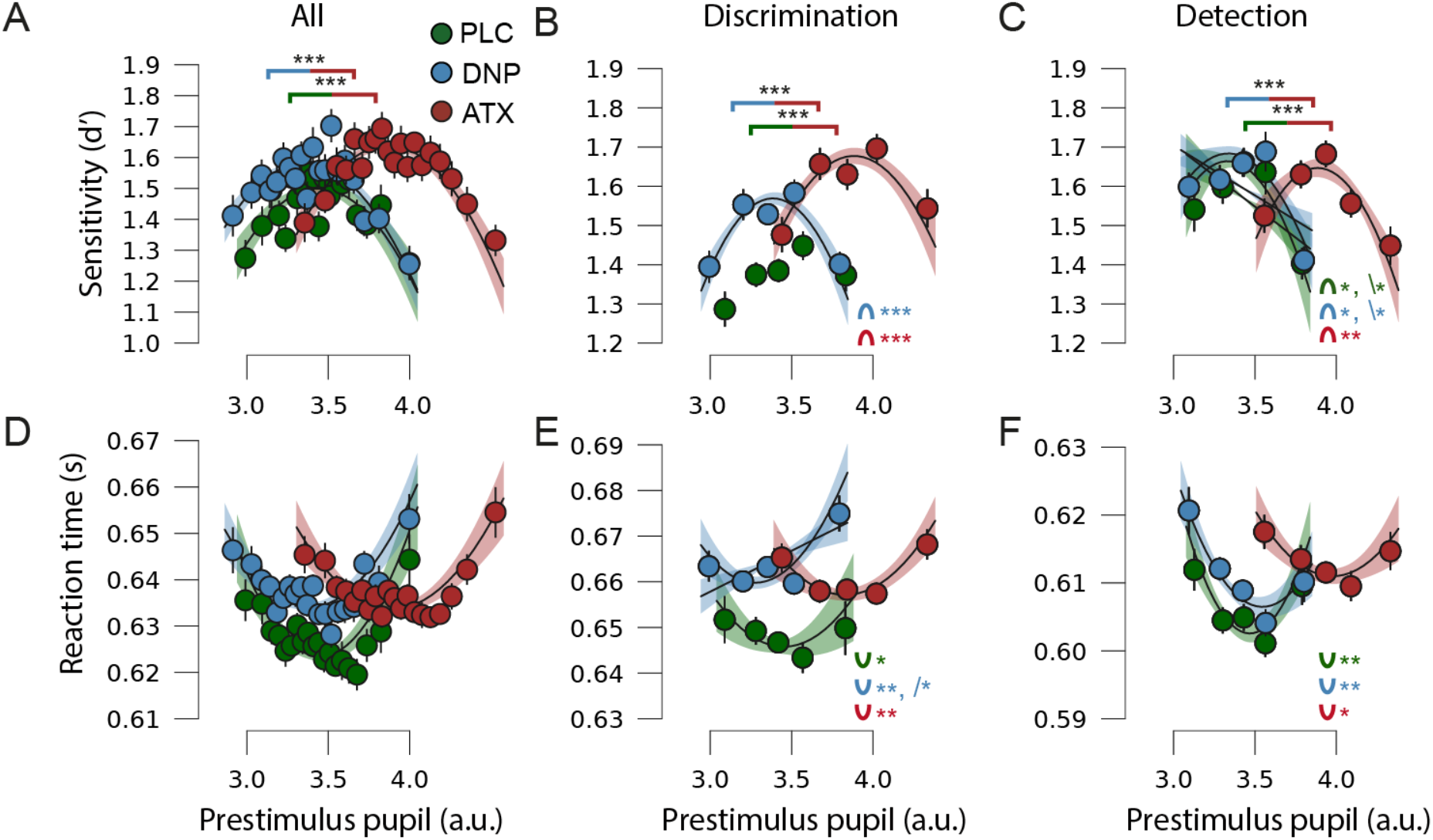
Arousal-performance curves. (A) Perceptual sensitivity (d’) for small to large prestimulus pupils for all tasks combined (20 bins), (B) for discrimination tasks (5 bins) and (C) for detection tasks split for Placebo (PLC), Donepezil (DNP), and Atomoxetine (ATX). (D) reaction time (RT) for small to large prestimulus pupils for all tasks combined (20 bins), (E) for discrimination tasks (5 bins) and (F) for detection tasks. Polynomial regression lines for first and second order fits (only significant fits are shown; significance is indicated in the bottom right of each panel, shading reflects SEM). Error bars on the data points represent SEM of the within-subject variation across tasks. Brackets above the curves indicate significant pupil size differences between drug conditions, indicating the rightward shift for ATX vs. the rest. ***p<.001, **p<.01, *p<.05, means and statistics in **Supplements**.

If ACh or catecholamines play a significant role in the modulation of performance, we expect DNP or ATX to change the shape and/or position of the pupil-linked arousal-performance relationship. In the DNP condition, for which pupil-linked arousal seemed unmodulated (i.e., position on the arousal spectrum was unaltered), the shape of the arousal-performance curves appeared unaffected and probabilistic model comparison favored the quadratic model for d’ and RT (all ΔAIC>30.1, all ΔBIC>25.8; **Figure 2A and D**, blue dots; **Supplementary table 2**). For ATX, ΔAIC and ΔBIC were also strongly in favor of the quadratic model for both sensitivity and RT in the ATX condition (all ΔAIC>36.6, all ΔBIC>32.2; **Figure 2A and D**, red dots; see also **Supplementary table 2**). Strikingly, the shape of the arousal-performance relationship seemed unaffected under ATX even though the position of the entire quadratic curve was now shifted to the right due to the overall increase in pupil size as compared to PLC (and other physiological arousal measures; indicative of a heightened arousal state; **Figure 1E-F**).

Our previous work indicated that the inverted U-shaped pupil-performance relationship does not depend on decision type (detection vs. discrimination) under neutral circumstances^1^. Yet, because the pharmacological manipulation may have affected discrimination and detection decisions differently, we split up our data according to decision type. To accommodate to the lowered statistical power after splitting the data, we now used five pupil bins combined with polynomial regression, identical to our previous work^1^. Again, the pupil-performance relationship was overall quadratically shaped for both sensitivity and RT for all drugs and decision-types (**Figure 2B-C and 2E-F**), although there was also evidence for linear relationships in two cases (likely due to lower trial counts and fewer bins used for fits, details in **Supplements**). As can be seen in **Figure 2B**, besides an overall rightward shift of the arousal-performance curve for ATX, ATX also selectively improved overall sensitivity on the discrimination tasks as compared to PLC (t(27)=2.42, p=.02), reflected in an additional upward shift of the ATX arousal-performance curve. ATX did not influence sensitivity on the detection tasks (t(27)=0.06, p=.95), nor did it influence RTs on either of the tasks (both ps>.27, more statistics in **Supplements**). DNP did not affect mean sensitivity or RTs for either decision type (all ps>.17; see **Supplements**). In sum, ATX administration appeared to affect discrimination decisions differently, but it did not change the general shape of the pupil-performance relationship.

In all conditions, the intact pupil-performance curve under ATX was shifted rightward on the pupil size axis in line with the relative arousal scenario (**Figure 1B**). Given that catecholamines are extensively linked to Yerkes-Dodson-like relationships^15,26,29^, that pharmacological manipulation of catecholamines has been shown to affect behavioral decision-making (i.e., **Figure 2B** and ^26,28–32)^, and that we observed a clear increase of the arousal state of our participants after ingestion of ATX, including increases in heart rate and blood pressure (**Figure 1H-I**), it seems likely that catecholamines are involved in maintaining the inverted U-shaped arousal-performance relationship, but apparently not in the most straightforward manner. To mechanistically explain how ATX could shift the arousal-performance curve on the arousal axis, essentially normalizing performance within each arousal state, we extended a neurobiologically plausible model that we have previously developed^1^. With this model, we have demonstrated that the Yerkes-Dodson curve can be achieved by the influence of arousal fluctuations on two types of interneurons (vasoactive intestinal peptide; VIP, and somatostatin; SST) that together form a disinhibitory pathway for the excitatory neural populations that encode the available sensory evidence (E_A_ and E_B_ encoding the evidence for choice A and B; **Figure 3A**; details in Methods). The model delineates a cortical circuit performing a detection and/or discrimination task under the influence of an arousal signal that is highly correlated to pupil size. It furthermore incorporates a non-selective population of parvalbumin (PV) interneurons, whose role is to provide a baseline level of inhibition and to mediate the competition between excitatory populations E_A_ and E_B_. Both excitatory populations receive sensory input, prompting them to increase their own activity and suppress the activity of the other excitatory population –a process mediated by the inhibitory population PV. A decision is made when the firing rate of one of the excitatory populations reaches a certain predefined threshold, giving rise to a winner-take-all decision process^47^, for a detection (A: present, B: absent) or a discrimination decision (choice A, choice B).

**Figure 3.**
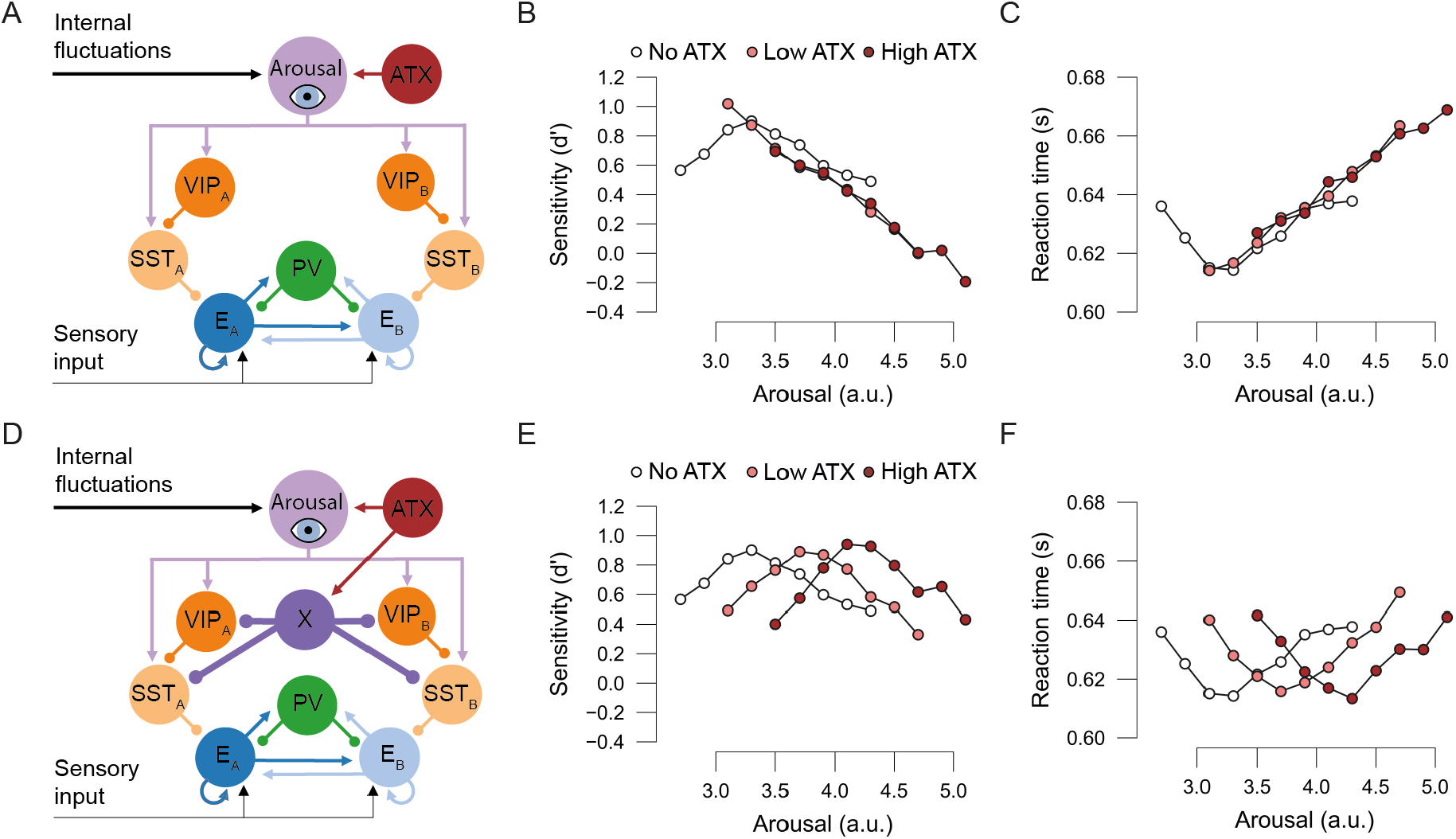
Computational modelling. (A) Schematic description of the first model. Arousal is determined by the combination of arousal fluctuations and the ATX signal (no/low/high ATX). Excitatory populations E_A_ and E_B_ encode two possible choices in a decision-making task (choice A and B). E_A_ and E_B_ are modulated by the arousal signal to VIP and SST cells. PV interneurons mediate the competition between E_A_ and E_B_. Connectivity follows electrophysiological evidence and previous modeling approaches. Lines with arrows/dots indicate excitatory/inhibitory connections, respectively. (B) Arousal-performance curves derived from the first model, expressed in d’ and RT (C) as a function of arousal plotted for no/low/high ATX. (D) Schematic description of the second model. ATX increases arousal, but also excites neural population X, which subsequently inhibits the inhibitory chain of VIP and SST cells. (E) Arousal-performance curves of the second model for d’and RT (F).

We extended the model with a neuromodulation (ATX) signal to gain a better understanding of the mechanism behind the observed shift of the arousal-performance curve after ingestion of ATX. As a first step, we implemented what we considered to be the simplest solution: an ATX signal that selectively increases arousal (**Figure 3A**) in line with the influence of ATX on physiological arousal measures that we observed (**Figure 1E-I**). Note that the model now has two “arousal inputs” that together determine arousal: arousal fluctuations as well as ATX induced arousal (state) enhancement. These two arousal inputs are summed into the arousal node (lilac in **Figure 3A**), whose output is plotted on the x-axis of our modelling results figures, as it directly translates to the arousal state that we measure with pupil size in our experimental studies.

When we simulated performance of this version of the model with two levels of ATX (low/high ATX) and compared this to baseline (no ATX), we observed that the relationship between performance measures and arousal became linear whilst shifting on the arousal axis (**Figure 3B** and **3C**), and this implementation of the model was therefore not able to mimic our findings. We thus considered a second, more elaborate model architecture in which ATX enhances arousal, but simultaneously influences other brain areas, which may have additional modulatory effects on neural circuits involved in the task. More concretely, we introduced a non-specific catecholamine-selective population that is excited by ATX but downregulates activity of both VIP and SST neurons (region “X”, **Figure 3D**). Note that we assume that the (negative) modulation strength from the catecholamine-selective area X to VIP and SST is stronger than the (positive) modulation strength of summed arousal to population VIP and SST (i.e., thicker purple lines from X to VIP/SST in **Figure 3B**), otherwise the opposing effects of these connections would cancel each other out or be overcompensated. This new model nicely echoes our experimental observations. First, when ATX increases above baseline (no ATX/PLC; **Figure 3E-F**) levels, arousal is increased as observed experimentally in pupil size. Second, the firing of SST and VIP cells is weakly excited via the increased arousal level, but strongly inhibited by the catecholamine-selective population X that is activated by ATX. This means that the working point of the VIP-SST-pyramidal cell disinhibitory pathway is altered, providing more inhibition to the excitatory populations than under no ATX/PLC. In other words, the net effect of (low/high) ATX is stronger inhibition of the excitatory populations as compared to no ATX/PLC. As a result, the shape of the arousal-performance curves will remain intact but peak performance under ATX will occur at higher levels of arousal as compared to PLC. With the administration of ATX, the arousal-performance curve thus shifts to the right on the arousal (i.e. pupil size) axis, and more ATX equals a larger shift (**Figure 3C-D**). Taken together, we present a new model in which ATX causes a rightward shift of the arousal-performance relationship via a population of catecholamine-selective neurons that inhibit VIP and SST populations. In the discussion section we will elaborate on which brain region (or neural population) may fulfill the role of node X in our model.

## Discussion

Here we demonstrated that the relationship between pupil-linked arousal and task performance is inverted U-shaped, even when the arousal state is heightened due to the administration of the catecholaminergic drug atomoxetine (ATX). Strikingly, ATX shifted the arousal-performance curve rightward on the (pupil-linked) arousal axis without altering its overall shape. This suggests that task performance is not simply tied to the absolute level of arousal, but it is rather optimized relative to fluctuations within a given arousal state, supporting the relative arousal scenario (**Fig. 1B**). The relationship between arousal and performance may thus be more adaptive than previously understood, potentially operating through mechanisms similar to well-known normalization processes in neuroscience^33–35^. Arousal typically fluctuates around a midpoint, producing a normal distribution of arousal fluctuations within each arousal state, even in high arousal states (i.e., red curve in **Figure 1E** and see^5,6^). By centering the arousal-performance curve around this midpoint, organisms optimize performance for the most frequently encountered arousal levels within each state, ensuring overall higher performance compared to a flat or linear relationship. Previous research has indeed demonstrated that organisms can adapt their overall arousal state to maximize performance^6,48,49^, in line with the adaptive gain theory^15^. For example, mice adjust their arousal state to an optimal level in high-utility task contexts, partially by suppressing irrelevant locomotor behavior^6^. Together, these findings underscore the flexibility and efficiency of neural circuits responsible for shaping the arousal-performance relationship, both within and across arousal states.

At first glance, our findings may seem at odds with the common interpretation of the Yerkes-Dodson law, which states that performance should always peak during states of (global) moderate arousal. There is indeed evidence in animals, humans and computational modeling that performance is impaired during very low (i.e., drowsiness) or very high (i.e., locomotion) arousal states as compared to neutral states^5,6,9,46,50–53^. Yet, this body of work often reduces low and high arousal states to one or two data points, neglecting the shape of the arousal-performance relationship *within* these states, or more subtle arousal fluctuations in general. The Yerkes-Dodson law and the relative arousal scenario can coexist, as (relative) arousal fluctuations may shape performance *within* states on top of overall performance differences that occur *between* arousal states.

An important, yet open question is how relative arousal shapes performance over time. In our study, we manipulated arousal state across sessions, on different days, allowing participants to establish a new equilibrium for optimal performance in each session. At present, it is unclear how long an organism must spend in a new arousal state for the neural circuitry to adapt, and thus for the inverted U-shaped arousal-performance relationship to shift. Future experiments could compare the arousal-performance relationship during session-wise arousal state manipulations, as in our experiment, to block-wise manipulations (e.g., alternating blocks of stationarity and locomotion). This approach would help determine if the inverted U-shaped arousal-performance relationship requires extended time in a state to emerge.

Possibly surprisingly, ATX selectively improved mean task performance for the discrimination tasks compared to PLC, whilst the Yerkes-Dodson law would predict the opposite. However, with only two arousal states (PLC and ATX) one cannot establish the relative positions of those states on the Yerkes-Dodson curve. It is notoriously difficult to establish the (starting) point of individuals on the Yerkes-Dodson-curve and some auxiliary measures have been proposed as proxies (e.g., working memory capacity)^26,29,54^, but so far not satisfactorily. The observed data pattern suggests that participants were slightly under-aroused during the PLC condition, which is common during long experiments, and closer to peak performance in the ATX condition.

In contrast to ATX, DNP did not appear to modulate arousal. It is not uncommon that DNP does not elicit any changes in physiological arousal, even in the presence of behavioral or neural effects^25,27,31,32,37–39^, which we did not observe either. Our results do not allow us to dissociate between two possible conclusions about the role of acetylcholine: 1) either the cholinergic manipulation was ineffective, or 2) DNP was effective, but acetylcholine does not disrupt the shape of the arousal-performance relationship. Future work with a higher dose of DNP, other cholinergic agents (e.g., galantamine, rivastigmine), or optogenetics could provide more insight into the role of acetylcholine in the arousal-performance relationship (for more discussion see ^8,31,32^).

To provide a potential neural mechanism that explains our observations, we extended a computational model that we posited in previous work in two ways^1^: 1) we added a neuromodulation signal that heightened (summed) arousal, and 2) we included an additional brain region (called X), which strongly inhibits the activity of SST and VIP cells. There are several lines of evidence suggesting that prefrontal cortex (PFC) could potentially fulfill the role of region X in our model. First, acute administration of ATX mainly increases extracellular levels of noradrenaline and dopamine in PFC, and leaves extracellular catecholamine levels in other catecholamine-responsive regions (e.g., striatum and nucleus accumbens) unaffected in rodents^55,56^. Second, PFC, and in particular the orbitofrontal (OFC) and anterior cingulate cortex (ACC), plays an important role in the evaluation of task utility^15,57^. More specifically, these regions monitor the costs and benefits of task performance, which is essential for adapting neuromodulation levels to optimize performance. Moreover, OFC and ACC send strong convergent projections to the noradrenergic locus coeruleus (LC) in monkeys^11,15,58^. This connectivity allows LC to adapt neuromodulation (and consequently arousal) levels to task utility information that comes from high-level structures. PFC may direct similar information about task utility to the disinhibitory chain of VIP/SST interneurons, analogous to the role of population X in our model. Indeed, top-down projections from frontal/motor cortices are known to target SST and VIP cells in rodents leading to disinhibitory effects in sensory areas^59^. Such top-down modulation to SST/VIP is dependent on behavioral strategies^60^, again suggesting the involvement of high association areas like PFC. By modulating the activity of these interneurons, PFC can effectively influence the arousal-performance relationship, ensuring that performance remains optimal within different arousal states.

The computational model presented here considers inhibition and disinhibition as fundamental ingredients for the effect of arousal on performance. Besides its importance in the Yerkes-Dodson law^1^, inhibition has been highlighted as a key component in arousal-mediated regulation of visual processing^61,62^ and normalization of activity in attention models in vision^33,34^. Mapping normalization models to concrete biophysical implementations is a difficult and unresolved problem, although subtractive, divisive and nonlinear transformations needed for normalization effects to occur, may emerge from a combination of inhibition and noise^63^, both existing ingredients of our model. Exploring our proposed mechanism in more detailed models including PV, SST and VIP cells^64–66^ should shed more light onto the particular circuits and layers involved in arousal-regulated perception.

## Methods

### Subjects

30 right-handed Dutch speaking male participants (aged between 18-30) were recruited from the University of Amsterdam for this study. Because this study involved a pharmacological manipulation, all participants underwent extensive medical and psychological screening to rule out any medical or mental illnesses. All participants gave written consent for participation and received monetary compensation. This study was approved both by the local ethics committee of the University of Amsterdam and the Medical Ethical Committee of the Amsterdam Medical Centre. All participants provided written informed consent after explanation of the experimental protocol. Two participants decided to withdraw from the experiment after the first experimental session. The data from these participants was excluded from further analyses, resulting in N=28 for the results presented here.

### Screening procedure

After completing the registration process, potential participants were contacted via e-mail to communicate the inclusion criteria, specific procedures, and potential risks associated with the study. Additionally, they were provided with the contact information of an independent physician whom they could reach out to for any inquiries or further clarifications. Following a contemplation period of seven days, candidates were contacted again to invite them for a preliminary phone discussion. This pre-intake conversation aimed to verify that the candidate indeed fulfilled all the necessary inclusion and exclusion criteria (a comprehensive list of criteria can be found in the **Supplements**). If the requirements were met, candidates were then invited to participate in an on-site intake session at the research facility of the University of Amsterdam. During this face-to-face intake session, the experimental protocol was thoroughly explained, including the subsequent physical and mental evaluations, after which candidate participants were asked to provide their written consent. The intake process also encompassed a range of physiological measurements such as body mass index (BMI), heart rate, blood pressure (BP), and an electrocardiogram (ECG). Additionally, a psychiatric questionnaire was administered to assess the candidates’ mental well-being. The information gathered during the intake was carefully reviewed by a physician, who subsequently determined the eligibility of each candidate participant. Finally, participants performed the staircasing procedure for the behavioral tasks (for more details, see **Staircasing procedure**).

### Drug administration

This study employed a randomized, double-blind crossover design. Atomoxetine (ATX; 40mg), donepezil (DNP; 5mg) and placebo (PLC) were administered on different experimental sessions, all separated by a minimum of seven days. The sequence in which the drugs were given was counterbalances among the participants. The experimental days commenced at 9am and concluded at 4pm. Since ATX (∼2 hours) and DNP (∼4 hours) reach their peak plasma levels at distinct times, they were administered prior to the initiation of the behavioral tasks at different intervals (**Figure 1C**). To maintain the integrity of the double-blind setup, participants were instructed to orally consume a pill four hours before the commencement of the initial behavioral task. This pill could either contain DNP or PLC. Subsequently, a second pill was ingested two hours prior to the start of the same task, which could contain either ATX or PLC. This ensured that participants received either PLC along with an active pharmaceutical or two PLCs during each experimental session. Additional details regarding ATX and DNP can be found in the **Supplements**.

### Procedure

Participants performed multiple tasks during a single experimental session. During the first two hours of each session, participants performed auditory discrimination and detection tasks that are not part of this study. In the second half of all sessions, participants performed four versions of visual detection and discrimination tasks. The order of the tasks was randomized between participants, but constant over sessions for each participant.

Below, we provide a brief description of the behavioral tasks, for additional results please see^31^. During all tasks, participants were seated 80cm from a computer monitor (69×39cm, 60Hz, 1920×1080 pixels) in a darkened, sound isolated room. To minimize head movements, participants rested their heads on a head-mount with chinrest. All tasks were programmed in Python 2.7 using PsychoPy^67^ and in-house scripts.

### Behavioral tasks

Participants performed four visual tasks. Two of these were variations of a discrimination task, in which participants were asked to discriminate the orientation of a Gabor patch hidden in dynamic visual noise as being rotated clockwise (CW: 45°, 50% of trials) or counterclockwise (CCW: −45°, 50% of trials; **Figure 1D**). The other two tasks were visual detection tasks in which participants had to indicate whether they believed a Gabor patch (CCW or CW) to be present or absent (50% present trials). The two detection tasks differed in terms of response bias manipulation (conservative versus liberal, see below). Besides these main task characteristics, there were some additional instructions and manipulations that are discussed below. During all tasks, the online measurement of gaze position ensured that participants maintained fixation. Trials in which gaze deviated by more than 1.5° from fixation on the horizontal axis were excluded from subsequent analyses.

### Cued visual discrimination task

The first visual discrimination task was an adaptation of the Posner cueing task^68^. Target stimuli were presented unilaterally for a duration of 200ms, appearing either on the left or right side of the screen in equal proportions (50% left, 50% right). 1300ms prior to target presentation, a spatial cue that was predictive of the target stimulus location was presented for 300ms (80% cue validity). Participants were instructed to use this spatial cue to covertly shift their attention towards the cued location. Participants had the opportunity to respond within 1400 milliseconds after the onset of the target stimulus by pressing one of two buttons on the keyboard (*S* for CCW Gabor patches, *K* for CW). A variable inter-trial interval (ITI) randomly drawn from a uniform distribution between 250ms and 350ms started directly after a response or the end of the response window if no response was given. Participants performed 560 trials per session of this task, distributed over two runs of 280 trials.

### Uncued visual discrimination task

The other discrimination task took the form of a classical visual discrimination task. Target stimuli were presented centrally for a duration of 200 milliseconds without any additional manipulations. In addition to identifying the orientation of the target stimulus, participants were also required to provide their level of confidence in their decision. Prior to engaging in the task, participants received instructions to evenly distribute their confidence reports, a measure taken to ensure a balanced reporting of low and high confidence answers and prevent bias towards low confidence responses in this challenging task. Participants conveyed their orientation judgment and confidence simultaneously by pressing one of four designated buttons on the keyboard (A for high confidence CCW Gabor patches, S for low confidence CCW, K for low confidence CW, L for high confidence CW). Once again, participants had a time window of 1400 milliseconds to make their response, followed by a 250-350ms ITI. Participants completed a total of 600 trials for the uncued visual discrimination task, divided across two sets of 300 trials each.

### Liberal visual detection task

In the liberal visual detection task, target stimuli were presented centrally for 200ms as well. On every trial, the noise stimulus (a circle containing dynamic noise) was presented, while the target stimuli (Gabor patches) were only exhibited in 50% of the trials. Participants were instructed to determine whether they perceived a target stimulus or not, and they indicated their response by pressing *S* (for target absent) or *K* (for target present). These target stimuli retained the same CCW and CW orientations as in the discrimination tasks, but the orientation was not task-relevant and could be disregarded. The response window was again 1400ms from stimulus onset, followed by a 250-350ms ITI. Response bias was manipulated towards more liberal answers by means of negative auditory feedback in the form of a buzzer sound after missed target stimuli (i.e., misses in signal detection theory), presented immediately after the response. Participants performed 480 trials of the liberal visual detection task on each session, distributed over two runs of 240 trials.

### Conservative visual detection task

The conservative detection task was like the liberal detection task, differing solely in the manipulation of response bias towards more conservative responses. In this case, negative auditory feedback was provided after falsely reporting the presence of a target stimulus (i.e., false alarms in signal detection theory). Again, participants performed 480 trials (two runs of 240 trials) per session of this task.

### Staircasing procedure

We titrated performance on all tasks to 75% correct in a separate intake session that preceded the three experimental sessions discussed here^31^. To this end, the opacity of the Gabor patches (i.e., signal strength) was varied according to the weighted transformed up/down method proposed by Kaernbach^69^, whilst the visual noise was kept constant.

### Eye-tracking acquisition and preprocessing

Gaze position and pupil size were recorded with an EyeLink 1000 eye tracker (SR Research, Canada) during the experiment at 500Hz. Nine-point calibration was performed at the start of each run to ensure high data quality. Additionally, a head-mount with a chinrest was employed to minimize any potential head movements from participants. Participants were given explicit instructions to keep head movements to a minimum and to try to avoid blinking during trials.

The pupillometry data of all tasks was preprocessed in the exact same way. Pupil traces were lowpass filtered at 10Hz, blinks were linearly interpolated and the effects of blinks and saccades on pupil diameter were removed via finite impulse-response deconvolution^70^. Note that EyeLink provides pupil size in arbitrary units, which we divided by 1000 for plotting purposes.

## Data analysis

### Analysis for data collapsed across tasks

To assess the overall shape of the relationship between prestimulus pupil-linked arousal and perceptual decision-making, we first combined the data of all tasks, split for the pharmacological manipulation. To quantify prestimulus pupil-linked arousal, we took the minimally preprocessed pupil traces of all tasks, and we calculated the average pupil size in the baseline window (−500ms – 0ms) before each stimulus onset (**Figure 1J**). We excluded all trials for which the eyes were closed during the entire baseline window, as well as trials for which pupil size during the baseline window was smaller or larger than three standard deviations from a subjects’ mean baseline pupil size. Next, for each run of each task separately, we assigned each trial to one of twenty equally populated bins based on the average prestimulus pupil size (note that we use five bins for the <u>Analyses performed</u> <u>separately for decision type</u>). The binning procedure was performed per run because it is not possible to assess whether pupil size differences between individual runs are the result of arousal fluctuations or small shifts in the exact head location (even though we used a head-mount). Therefore, we are looking at arousal fluctuations that occur within experimental runs (**Figure 1K**, shown for five bins).

After having assigned all trials to twenty bins, we calculated Signal Detection Theoretic sensitivity (SDTs d’^41^) and average reaction time (RT) as our dependent measures of perceptual decision-making for each bin. We also calculated the mean pupil size for each bin to replace (equally spaced) bin numbers with (not necessarily equally spaced) mean prestimulus pupil size values, to do justice to the true relationship between pupil size and decision behavior.

Next, we subsequently averaged over runs and tasks, leaving us with mean d’ and RT for 20 bins for each subject and for each drug. To assess the shape of the relationship between arousal and perceptual decision-making, we next used mixed linear models (see below <u>Mixed</u> <u>linear models</u>).

### Mixed linear models

We used a mixed linear modeling approach implemented in the Python-package *Statsmodels*^71^ to quantify the dependence of behavioral sensitivity and reaction time on pupil size^72^. Specifically, we fitted two mixed models to test whether pupil response bin predominantly exhibited a monotonic effect (first-order), or a non-monotonic effect (second-order) on the behavioral metric of interest (*y*). The fixed effects were specified as:

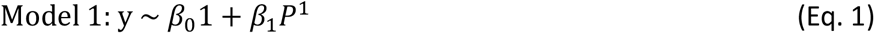

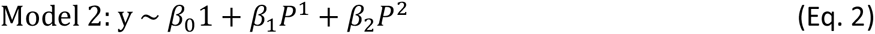

with *β* as regression coefficients and *P* as the average baseline pupil size in each bin. We included the maximal random effects structure justified by the design^73^: intercepts and pupil size coefficients could vary with participant. The mixed models were fitted through restricted maximum likelihood estimation. The two models were then formally compared based on Akaike Information Criterion (AIC)^42^ and Bayesian Information Criterion (BIC)^43^.

### Analyses performed separately for decision type

After having assessed that the shape of the overall relationship between pupil-linked arousal and perceptual decision-making was quadratically shaped for all drugs separately, we investigated whether this relationship also held for the different decision types in our dataset. We collapsed the data of all tasks over the relevant features (i.e., detection (2 tasks), discrimination (2 tasks)), but kept the data split for the pharmacological manipulations. We next treated the data as before (see <u>Analysis for data collapsed across all tasks</u>), but this time we assigned the trials to five equally populated bins (instead of twenty) to compensate for the lowered power after splitting the data (**Figure 1J-K**). Next, we performed polynomial regression (see below <u>Polynomial regression</u>) to assess the shape of the relationship between arousal and perceptual decision-making in the different decision types for each drug separately in our dataset.

### Polynomial regression

To assess the shape of the relationship between arousal and perceptual decision-making (quadratic or linear), we performed second order polynomial regression. Because we were merely interested in the shape of the relationship, and for visualization purposes, we first normalized our data in the pupil dimension (i.e., essentially centering the data around a common mean). Next, we modeled the relationship between our observed behavior (d’ and RT) and prestimulus pupil size as a negative quadratic relationship, and as a (unsigned) linear relationship using ordinary least squares linear regression with freely varying intercepts. We extracted the relevant beta coefficient from each model (ß_1_ for the linear model and ß_2_ for the quadratic model) for each subject and tested whether the coefficients were significantly different from zero using one-sample t-tests (α=.05). We first tested the significance of the linear model (two-sided), followed by the quadratic model. After having established that the overall relationship between prestimulus pupil and sensitivity/RT was negatively/positively quadratically shaped, respectively, we performed one-sided tests for the quadratic beta coefficients.

### Methods: Computational model

Following and extending our previous efforts, we considered a population-based model of neural dynamics to describe a general decision-making task^47^, which we adapted for our detection/discrimination tasks^1^. The model simulates, in a first instance, the temporal evolution of global synaptic conductance variables corresponding to the NMDA and GABA receptors of two competing excitatory populations and one inhibitory (PV) population. The model is described by the following equations:

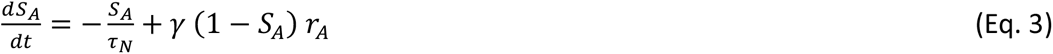

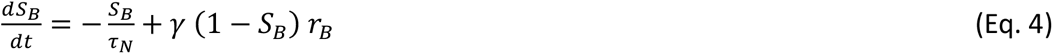

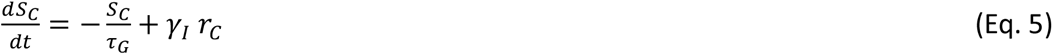

Above, S_A_ and S_B_ correspond, respectively, to the NMDA conductances of selective excitatory populations A and B, and S_C_ corresponds to the GABAergic conductance of the inhibitory population. The parameters in these equations take the following values: τ_N_=60 ms, τ_G_=5 ms, γ=1.282 and γ_I_=2. The variables r_A_, r_B_ and r_C_ are the mean firing rates of the two excitatory populations and one inhibitory population, respectively. We obtain their values by solving, at each time step, the transcendental equation *r_i_*= ϕ*_i_*(*I_i_*), with ϕ being a transfer function of the population (specified below) and I_i_ being the input to population ‘i’, given by:

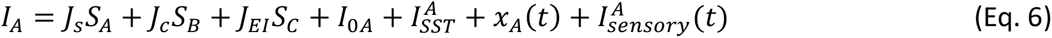

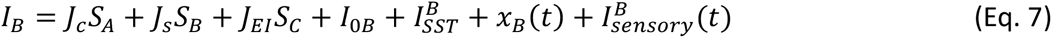

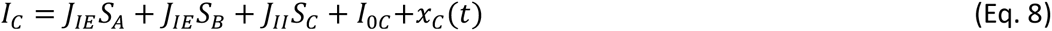

The parameters J_s_, J_c_ are the self- and cross-coupling synaptic terms between excitatory populations, respectively. J_EI_ is the coupling from the inhibitory populations to any of the excitatory ones, J_IE_ is the coupling from any of the excitatory populations to the inhibitory one, and J_II_ is the self-coupling strength of the inhibitory population. The parameters I_0i_ with i=A, B, C are background inputs to each population. Parameters in these equations take the following values: J_s_=0.49 nA, J_c_=0.0107 nA, J_IE_=0.3597 nA, J_EI_=-0.31 nA, J_II_=-0.12 nA, I_0A_=I_0B_=0.3294 nA and I_0C_=0.26 nA. The term I^i^_SST_ denotes the input to each excitatory population from its corresponding SST population (see details about this term below).

The term x_i_(t) with i=A, B, C is an Ornstein-Uhlenbeck process, which introduces some level of stochasticity in the system. It is given by:

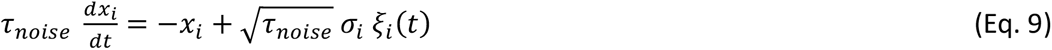

Here, ξ_i_(t) is a Gaussian white noise, the time constant is τ_noise_=2 ms and the noise strength is σ_A,B_=0.03 nA for excitatory populations and σ_C_=0 for the inhibitory one.

The last term in Eqs. 6 and 7 represents the external sensory input arriving to both populations. Assuming a detection task in which the subject has to detect the sensory stimulus A, the input is given by:

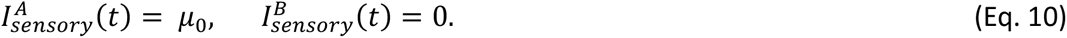

The average stimulus strength is μ_0_=0.0133 unless specified otherwise. The stimulus is present during the whole duration of the trial. The discrimination task can be simply simulated by alternating the population which receives the input in the equation above, and therefore provides equivalent results.

The transfer function ϕ_i_(t) which transform the input into firing rates takes the following form for the excitatory populations:

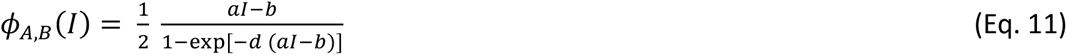

The values for the parameters are *a*=135 Hz/nA, *b*=54 Hz and *d*=0.308 s. For the inhibitory population a similar function can be used, but for convenience we choose a threshold-linear function:

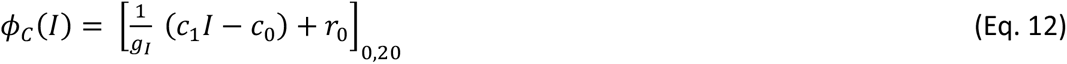

The notation [*x*]_0,20_ denotes a minimum value of zero (rectification) and a maximum value at 20 spikes/s (saturation). The values for the parameters are g_I_=4, c_1_=615 Hz/nA, c_0_=177 Hz and r_0_=5.5 spikes/s. It is sometimes useful for simulations (although not a requirement) to replace the transcendental equation *r_i_* = ϕ*_i_*(*I_i_*) by its analogous differential equation, of the form:

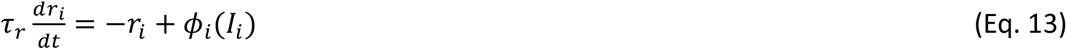

The time constant can take a typical value of τ_r_=2 ms.

In addition to the two selective excitatory populations and the PV population, our model also includes the effects of top-down input mediated by selective VIP and SST populations. To introduce this, we assume that the firing rate activity of VIP and SST cells is determined, respectively, by the following equations:

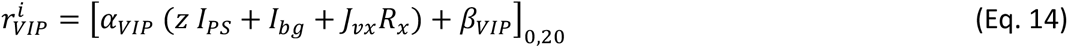

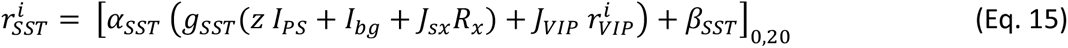

The subindex ‘i’ indicates the associated selective excitatory populations. Parameter values are α_VIP_=50, β_VIP_=0, α_SST_=20, α_SST_=32, g_SST_=2, J_VIP_=-0.1, z=0.1 and I_bg_=0.37. The variable R_x_ is the firing rate of unspecified, potentially neuromodulatory neurons (neural population X in **Figure 3D**) modulated by atomoxetine (ATX), which we assume to have an inhibitory effect over all SST and VIP populations (with synaptic couplings of J_sx_=-0.06 and J_vs_=-0.06, respectively) and follows the equation

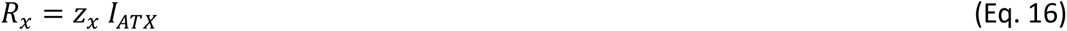

Here, z_x_=20 is a gain factor and I_ATX_ is the modulation of R_x_ by ATX, which is I_ATX_=0 for the control case, I_ATX_=0.05 nA for low ATX, and I_ATX_=0.1 nA for high ATX.

The parameter I_PS_ in Eqs. 12 and 13 indicates the variable associated with the pupil size, which is both influenced by the level of ATX and the underlying levels of arousal of the organism. It is given by:

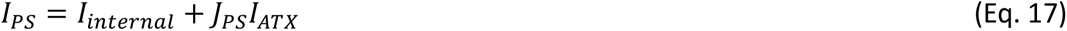

Here, we assume that the internally generated arousal levels are I_internal_=0.35 unless specified otherwise, and J_PS_=2 to capture the experimentally observed positive effect of ATX on pupil size.

Finally, we link the firing rate of the SST populations in Eq. 15 with the input to excitatory populations given in Eqs. 6 and 7 by assuming that:

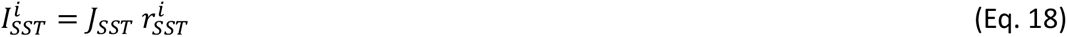

The synaptic strength is given by J_SST_=-0.001.

For each simulated trial, we consider that the circuit makes a detection when the firing rate of either the excitatory population A (or B, in the case of a discrimination task) reaches a threshold of 15 spikes/s. The duration of the trial is set to T_trial_ =1.5 sec. To compute sensitivity (d’) and reaction time, we averaged over 3000 trials for each value of the pupil size considered. In **Figure 3B-C** and **3D-E**, we plot performance for selected arousal ranges for each ATX level.

## Data, materials, and software availability

The data of all experiments, analysis scripts, as well as the computational modeling scripts can be found at https://osf.io/bczku/

## References

1. Beerendonk, L., Mejías, J.F., Nuiten, S.A., de Gee, J.W., Fahrenfort, J.J., and van Gaal, S. (2024). A disinhibitory circuit mechanism explains a general principle of peak performance during mid-level arousal. Proc. Natl. Acad. Sci. U. S. A. 121, e2312898121. 10.1073/pnas.2312898121.

2. Yerkes, R.M., and Dodson, J.D. (1908). The relation of strength of stimulus to rapidity of habit-formation. J. Comp. Neurol. Psychol. 18, 459–482.

3. Teigen, K.H. (1994). Yerkes-Dodson: A law for all seasons. Theory Psychol. 4, 525–547. 10.1177/0959354394044004.

4. Hebb, D.O. (1955). Drives and the C.N.S. (conceptual nervous system). Psychol. Rev. 62, 243–254. 10.1037/h0041823.

5. McGinley, M.J., David, S.V., and McCormick, D.A. (2015). Cortical Membrane Potential Signature of Optimal States for Sensory Signal Detection. Neuron 87, 179–192. 10.1016/j.neuron.2015.05.038.

6. de Gee, J.W., Mridha, Z., Hudson, M., Shi, Y., Ramsaywak, H., Smith, S., Karediya, N., Thompson, M., Jaspe, K., Jiang, H., et al. (2024). Strategic stabilization of arousal boosts sustained attention. Curr. Biol. 0. 10.1016/j.cub.2024.07.070.

7. Reimer, J., McGinley, M.J., Liu, Y., Rodenkirch, C., Wang, Q., McCormick, D.A., and Tolias, A.S. (2016). Pupil fluctuations track rapid changes in adrenergic and cholinergic activity in cortex. Nat. Commun. 7, 13289. 10.1038/ncomms13289.

8. Slater, C., Liu, Y., Weiss, E., Yu, K., and Wang, Q. (2022). The Neuromodulatory Role of the Noradrenergic and Cholinergic Systems and Their Interplay in Cognitive Functions: A Focused Review. Brain Sci 12. 10.3390/brainsci12070890.

9. Sabat, M., de Dampierre, C., and Tallon-Baudry, C. (2024). Evidence for domain-general arousal from semantic and neuroimaging meta-analyses reconciles opposing views on arousal. bioRxiv, 2024.05.27.594944. 10.1101/2024.05.27.594944.

10. Joshi, S., Li, Y., Kalwani, R.M., and Gold, J.I. (2016). Relationships between Pupil Diameter and Neuronal Activity in the Locus Coeruleus, Colliculi, and Cingulate Cortex. Neuron 89, 221–234. 10.1016/j.neuron.2015.11.028.

11. de Gee, J.W., Colizoli, O., Kloosterman, N.A., Knapen, T., Nieuwenhuis, S., and Donner, T.H. (2017). Dynamic modulation of decision biases by brainstem arousal systems. Elife 6. 10.7554/eLife.23232.

12. Murphy, P.R., O’Connell, R.G., O’Sullivan, M., Robertson, I.H., and Balsters, J.H. (2014). Pupil diameter covaries with BOLD activity in human locus coeruleus. Hum. Brain Mapp. 35, 4140–4154. 10.1002/hbm.22466.

13. Strauch, C., Wang, C.-A., Einhäuser, W., Van der Stigchel, S., and Naber, M. (2022). Pupillometry as an integrated readout of distinct attentional networks. Trends Neurosci. 45, 635–647. 10.1016/j.tins.2022.05.003.

14. Lloyd, B., de Voogd, L.D., Mäki-Marttunen, V., and Nieuwenhuis, S. (2023). Pupil size reflects activation of subcortical ascending arousal system nuclei during rest. Elife 12. 10.7554/eLife.84822.

15. Aston-Jones, G., and Cohen, J.D. (2005). An integrative theory of locus coeruleus-norepinephrine function: adaptive gain and optimal performance. Annu. Rev. Neurosci. 28, 403–450. 10.1146/annurev.neuro.28.061604.135709.

16. Waschke, L., Tune, S., and Obleser, J. (2019). Local cortical desynchronization and pupil-linked arousal differentially shape brain states for optimal sensory performance. Elife 8. 10.7554/eLife.51501.

17. Podvalny, E., King, L.E., and He, B.J. (2021). Spectral signature and behavioral consequence of spontaneous shifts of pupil-linked arousal in human. Elife 10. 10.7554/eLife.68265.

18. van Kempen, J., Loughnane, G.M., Newman, D.P., Kelly, S.P., Thiele, A., O’Connell, R.G., and Bellgrove, M.A. (2019). Behavioural and neural signatures of perceptual decision-making are modulated by pupil-linked arousal. Elife 8. 10.7554/eLife.42541.

19. Grujic, N., Tesmer, A., Bracey, E., Peleg-Raibstein, D., and Burdakov, D. (2023). Control and coding of pupil size by hypothalamic orexin neurons. Nat. Neurosci. 10.1038/s41593-023-01365-w.

20. de Lecea, L., Carter, M.E., and Adamantidis, A. (2012). Shining light on wakefulness and arousal. Biol. Psychiatry 71, 1046–1052. 10.1016/j.biopsych.2012.01.032.

21. Sander, C., Hensch, T., Wittekind, D.A., Böttger, D., and Hegerl, U. (2015). Assessment of wakefulness and brain arousal regulation in psychiatric research. Neuropsychobiology 72, 195–205. 10.1159/000439384.

22. Reimer, J., Froudarakis, E., Cadwell, C.R., Yatsenko, D., Denfield, G.H., and Tolias, A.S. (2014). Pupil fluctuations track fast switching of cortical states during quiet wakefulness. Neuron 84, 355–362. 10.1016/j.neuron.2014.09.033.

23. Rogers, S.L., and Friedhoff, L.T. (1998). Pharmacokinetic and pharmacodynamic profile of donepezil HCl following single oral doses. Br. J. Clin. Pharmacol. 46 Suppl 1, 1–6. 10.1046/j.1365-2125.1998.0460s1001.x.

24. Simpson, D., and Plosker, G.L. (2004). Atomoxetine: a review of its use in adults with attention deficit hyperactivity disorder. Drugs 64, 205–222. 10.2165/00003495-200464020-00005.

25. Boucart, M., Bubbico, G., Szaffarczyk, S., Defoort, S., Ponchel, A., Waucquier, N., Deplanque, D., Deguil, J., and Bordet, R. (2015). Donepezil increases contrast sensitivity for the detection of objects in scenes. Behav. Brain Res. 292, 443–447. 10.1016/j.bbr.2015.06.037.

26. Cools, R., and Arnsten, A.F.T. (2022). Neuromodulation of prefrontal cortex cognitive function in primates: the powerful roles of monoamines and acetylcholine. Neuropsychopharmacology 47, 309–328. 10.1038/s41386-021-01100-8.

27. Gratton, C., Yousef, S., Aarts, E., Wallace, D.L., D’Esposito, M., and Silver, M.A. (2017). Cholinergic, But Not Dopaminergic or Noradrenergic, Enhancement Sharpens Visual Spatial Perception in Humans. J. Neurosci. 37, 4405–4415. 10.1523/JNEUROSCI.2405-16.2017.

28. Gelbard-Sagiv, H., Magidov, E., Sharon, H., Hendler, T., and Nir, Y. (2018). Noradrenaline Modulates Visual Perception and Late Visually Evoked Activity. Curr. Biol. 28, 2239–2249.e6. 10.1016/j.cub.2018.05.051.

29. Cools, R., and D’Esposito, M. (2011). Inverted-U-shaped dopamine actions on human working memory and cognitive control. Biol. Psychiatry 69, e113–25. 10.1016/j.biopsych.2011.03.028.

30. Graf, H., Abler, B., Freudenmann, R., Beschoner, P., Schaeffeler, E., Spitzer, M., Schwab, M., and Grön, G. (2011). Neural correlates of error monitoring modulated by atomoxetine in healthy volunteers. Biol. Psychiatry 69, 890–897. 10.1016/j.biopsych.2010.10.018.

31. Nuiten, S.A., De Gee, J.W., Fahrenfort, J.J., and van Gaal, S. (2023). Catecholaminergic neuromodulation and selective attention jointly shape perceptual decision making. eLife. 10.7554/elife.87022.

32. Nuiten, S.A., De Gee, J.W., Zantvoord, J.B., Fahrenfort, J.J., and van Gaal, S. (2024). Pharmacological elevation of catecholamine levels improves perceptual decisions, but not metacognitive insight. eNeuro, ENEURO.0019–24.2024. 10.1523/ENEURO.0019-24.2024.

33. Reynolds, J.H., and Heeger, D.J. (2009). The normalization model of attention. Neuron 61, 168–185. 10.1016/j.neuron.2009.01.002.

34. Carandini, M., and Heeger, D.J. (2011). Normalization as a canonical neural computation. Nat. Rev. Neurosci. 13, 51–62. 10.1038/nrn3136.

35. Heeger, D.J. (2017). Theory of cortical function. Proc. Natl. Acad. Sci. U. S. A. 114, 1773– 1782. 10.1073/pnas.1619788114.

36. Diba, K., and Buzsáki, G. (2008). Hippocampal network dynamics constrain the time lag between pyramidal cells across modified environments. J. Neurosci. 28, 13448–13456. 10.1523/JNEUROSCI.3824-08.2008.

37. Pfeffer, T., Avramiea, A.-E., Nolte, G., Engel, A.K., Linkenkaer-Hansen, K., and Donner, T.H. (2018). Catecholamines alter the intrinsic variability of cortical population activity and perception. PLoS Biol. 16, e2003453. 10.1371/journal.pbio.2003453.

38. Pfeffer, T., Ponce-Alvarez, A., Tsetsos, K., Meindertsma, T., Gahnström, C.J., van den Brink, R.L., Nolte, G., Engel, A.K., Deco, G., and Donner, T.H. (2021). Circuit mechanisms for the chemical modulation of cortex-wide network interactions and behavioral variability. Science Advances 7, eabf5620. 10.1126/sciadv.abf5620.

39. Silver, M.A., Shenhav, A., and D’Esposito, M. (2008). Cholinergic enhancement reduces spatial spread of visual responses in human early visual cortex. Neuron 60, 904–914. 10.1016/j.neuron.2008.09.038.

40. Sheynin, Y., Rosa-Neto, P., Hess, R.F., and Vaucher, E. (2020). Cholinergic Modulation of Binocular Vision. J. Neurosci. 40, 5208–5213. 10.1523/JNEUROSCI.2484-19.2020.

41. Green, D.M., and Swets, J.A. (1966). Signal detection theory and psychophysics (Wiley New York).

42. Akaike, H. (1974). A new look at the statistical model identification. IEEE Trans. Automat. Contr. 19, 716–723. 10.1109/TAC.1974.1100705.

43. Schwarz, G. (1978). Estimating the dimension of a model. Ann. Stat. 6, 461–464. 10.1214/aos/1176344136.

44. Burnham, K.P., and Anderson, D.R. (2004). Multimodel inference. Sociol. Methods Res. 33, 261–304. 10.1177/0049124104268644.

45. Murphy, P.R., Robertson, I.H., Balsters, J.H., and O’connell, R.G. (2011). Pupillometry and P3 index the locus coeruleus-noradrenergic arousal function in humans. Psychophysiology 48, 1532–1543. 10.1111/j.1469-8986.2011.01226.x.

46. Neske, G.T., Nestvogel, D., Steffan, P.J., and McCormick, D.A. (2019). Distinct Waking States for Strong Evoked Responses in Primary Visual Cortex and Optimal Visual Detection Performance. J. Neurosci. 39, 10044–10059. 10.1523/JNEUROSCI.1226-18.2019.

47. Wong, K.-F., and Wang, X.-J. (2006). A recurrent network mechanism of time integration in perceptual decisions. J. Neurosci. 26, 1314–1328. 10.1523/JNEUROSCI.3733-05.2006.

48. Faller, J., Cummings, J., Saproo, S., and Sajda, P. (2019). Regulation of arousal via online neurofeedback improves human performance in a demanding sensory-motor task. Proc. Natl. Acad. Sci. U. S. A. 116, 6482–6490. 10.1073/pnas.1817207116.

49. Hulsey, D., Zumwalt, K., Mazzucato, L., McCormick, D.A., and Jaramillo, S. (2024). Decision-making dynamics are predicted by arousal and uninstructed movements. Cell Rep. 43, 113709. 10.1016/j.celrep.2024.113709.

50. Bullock, T., Elliott, J.C., Serences, J.T., and Giesbrecht, B. (2017). Acute Exercise Modulates Feature-selective Responses in Human Cortex. J. Cogn. Neurosci. 29, 605– 618. 10.1162/jocn_a_01082.

51. Beaman, C.B., Eagleman, S.L., and Dragoi, V. (2017). Sensory coding accuracy and perceptual performance are improved during the desynchronized cortical state. Nat. Commun. 8, 1308. 10.1038/s41467-017-01030-4.

52. Sörensen, L.K.A., Bohté, S.M., Slagter, H.A., and Scholte, H.S. (2022). Arousal state affects perceptual decision-making by modulating hierarchical sensory processing in a large-scale visual system model. PLoS Comput. Biol. 18, e1009976. 10.1371/journal.pcbi.1009976.

53. Shine, J.M., Müller, E.J., Munn, B., Cabral, J., Moran, R.J., and Breakspear, M. (2021). Computational models link cellular mechanisms of neuromodulation to large-scale neural dynamics. Nat. Neurosci. 24, 765–776. 10.1038/s41593-021-00824-6.

54. Cools, R., Gibbs, S.E., Miyakawa, A., Jagust, W., and D’Esposito, M. (2008). Working memory capacity predicts dopamine synthesis capacity in the human striatum. J. Neurosci. 28, 1208–1212. 10.1523/JNEUROSCI.4475-07.2008.

55. Koda, K., Ago, Y., Cong, Y., Kita, Y., Takuma, K., and Matsuda, T. (2010). Effects of acute and chronic administration of atomoxetine and methylphenidate on extracellular levels of noradrenaline, dopamine and serotonin in the prefrontal cortex and striatum of mice: Acute and chronic treatment with ADHD drugs. J. Neurochem. 114, 259–270. 10.1111/j.1471-4159.2010.06750.x.

56. Bymaster, F.P., Katner, J.S., Nelson, D.L., Hemrick-Luecke, S.K., Threlkeld, P.G., Heiligenstein, J.H., Morin, S.M., Gehlert, D.R., and Perry, K.W. (2002). Atomoxetine increases extracellular levels of norepinephrine and dopamine in prefrontal cortex of rat: a potential mechanism for efficacy in attention deficit/hyperactivity disorder. Neuropsychopharmacology 27, 699–711. 10.1016/S0893-133X(02)00346-9.

57. Botvinick, M.M., Braver, T.S., Barch, D.M., Carter, C.S., and Cohen, J.D. (2001). Conflict monitoring and cognitive control. Psychol. Rev. 108, 624–652. 10.1037/0033-295x.108.3.624.

58. Joshi, S., and Gold, J.I. (2022). Context-dependent relationships between locus coeruleus firing patterns and coordinated neural activity in the anterior cingulate cortex. Elife 11. 10.7554/eLife.63490.

59. Pfeffer, C.K., Xue, M., He, M., Huang, Z.J., and Scanziani, M. (2013). Inhibition of inhibition in visual cortex: the logic of connections between molecularly distinct interneurons. Nat. Neurosci. 16, 1068–1076. 10.1038/nn.3446.

60. Piet, A., Ponvert, N., Ollerenshaw, D., Garrett, M., Groblewski, P.A., Olsen, S., Koch, C., and Arkhipov, A. (2024). Behavioral strategy shapes activation of the Vip-Sst disinhibitory circuit in visual cortex. Neuron 112, 1876–1890.e4. 10.1016/j.neuron.2024.02.008.

61. Klaver, L.M.F., Brinkhof, L.P., Sikkens, T., Casado-Román, L., Williams, A.G., van Mourik-Donga, L., Mejías, J.F., Pennartz, C.M.A., and Bosman, C.A. (2023). Spontaneous variations in arousal modulate subsequent visual processing and local field potential dynamics in the ferret during quiet wakefulness. Cereb. Cortex 33, 7564–7581. 10.1093/cercor/bhad061.

62. Crombie, D., Spacek, M.A., Leibold, C., and Busse, L. (2024). Spiking activity in the visual thalamus is coupled to pupil dynamics across temporal scales. PLoS Biol. 22, e3002614. 10.1371/journal.pbio.3002614.

63. Mejias, J.F., Payeur, A., Selin, E., Maler, L., and Longtin, A. (2014). Subtractive, divisive and non-monotonic gain control in feedforward nets linearized by noise and delays. Front. Comput. Neurosci. 8, 19. 10.3389/fncom.2014.00019.

64. Moreni, G., Pennartz, C.M.A., and Mejias, J.F. (2023). Synaptic plasticity is required for oscillations in a V1 cortical column model with multiple interneuron types. bioRxiv, 2023.08.27.555009. 10.1101/2023.08.27.555009.

65. Jiang, H.-J., Qi, G., Duarte, R., Feldmeyer, D., and van Albada, S.J. (2023). A Layered Microcircuit Model of Somatosensory Cortex with Three Interneuron Types and Cell-Type-Specific Short-Term Plasticity. bioRxiv, 2023.10.26.563698. 10.1101/2023.10.26.563698.

66. Moreni, G., Pennartz, C.M.A., and Mejias, J.F. (2024). Cell type specific firing patterns in a V1 cortical column model depend on feedforward and feedback activity. bioRxiv, 2024.04.02.587673. 10.1101/2024.04.02.587673.

67. Peirce, J.W. (2007). PsychoPy--Psychophysics software in Python. J. Neurosci. Methods 162, 8–13. 10.1016/j.jneumeth.2006.11.017.

68. Posner, M.I. (1980). Orienting of attention. Q. J. Exp. Psychol. 32, 3–25. 10.1080/00335558008248231.

69. Kaernbach, C. (1991). Simple adaptive testing with the weighted up-down method. Percept. Psychophys. 49, 227–229. 10.3758/bf03214307.

70. Knapen, T., de Gee, J.W., Brascamp, J., Nuiten, S., Hoppenbrouwers, S., and Theeuwes, J. (2016). Cognitive and Ocular Factors Jointly Determine Pupil Responses under Equiluminance. PLoS One 11, e0155574. 10.1371/journal.pone.0155574.

71. Seabold, S., and Perktold, J. (2010). Statsmodels: Econometric and statistical modeling with python. In Proceedings of the 9th Python in Science Conference (SciPy). 10.25080/majora-92bf1922-011.

72. de Gee, J.W., Tsetsos, K., Schwabe, L., Urai, A.E., McCormick, D., McGinley, M.J., and Donner, T.H. (2020). Pupil-linked phasic arousal predicts a reduction of choice bias across species and decision domains. Elife 9. 10.7554/eLife.54014.

73. Barr, D.J., Levy, R., Scheepers, C., and Tily, H.J. (2013). Random effects structure for confirmatory hypothesis testing: Keep it maximal. J. Mem. Lang. 68. 10.1016/j.jml.2012.11.001.

