## Supplements for "Adaptive arousal regulation: Pharmacologically shifting the peak of the Yerkes-Dodson curve by catecholaminergic enhancement of arousal"

**Catecholaminergic, but not cholinergic enhancement upregulates physiological arousal**

To assess whether atomoxetine (ATX) and donepezil (DNP) upregulated arousal as intended, we analyzed various physiological arousal measures. To index arousal, we measured baseline pupil size (mean pupil dilation in the 500ms leading up to stimulus onset), pupil size variation, heart rate (HR) and mean arterial blood pressure (BP). As there was no significant difference between prestimulus pupil size during the tasks that took place before blood plasma concentrations of the drugs peaked (not part of this manuscript; $F_{2,50}$=0.76, p=.47), we here compare raw prestimulus pupil size between pharmacological conditions. We observed a main effect of drug on prestimulus pupil size ($F_{2,54}$=25.5, p<.001), as well as a main effect of task type on prestimulus pupil size ($F_{1,27}$=4.69, p=.04), but no significant interaction effect between drug and task type ($F_{2,54}$=2.95, p=.06, see **Supplementary Table 1** for means). Post-hoc t-tests indicated that prestimulus pupil size was enlarged by ATX (t(27)=4.78, p<.001; **Figure 1F**), but not by DNP (t(27)=-0.72, p=.48; **Figure 1F**), as compared to PLC, as evidenced by a clear right-ward shift of the prestimulus pupil size distribution (**Figure 1E**). In addition, prestimulus pupil size of the ATX condition was significantly larger than pupil size of the DNP condition (t(27)=6.48, p<.001).

As for the effect of task type, post-hoc t-tests showed that prestimulus pupil size was larger for detection tasks as compared to discrimination tasks (t(27)=2.29, p=.03). Within the discrimination tasks, prestimulus pupil size was significantly larger for ATX (t(27)=4.55, p<.001), but not by DNP (t(27)=-1.17, p=.25), as compared to PLC. Again, prestimulus pupil size was significantly larger for ATX than for DNP (t(27)=6.55, p<.001). Prestimulus pupil size differences between drugs within the detection tasks were very similar to the discrimination tasks. ATX significantly increased prestimulus pupil size as compared to PLC (t(27)=4.88, p<.001), but DNP did not (t(27)=-0.18, p=.86). Again, prestimulus pupil size was significantly larger for ATX as compared to DNP (t(27)=6.23, p<.001). Overall, ATX appeared to affect prestimulus pupil size for the detection and discrimination tasks in a similar fashion.

To further assess the influence of the pharmacological manipulation on pupil-linked arousal, we repeated these analyses with prestimulus pupil size variation as the dependent variable. In other words, we assessed whether the width of the prestimulus pupil size distribution changed after ingestion of ATX or DNP. We observed a significant main effect of drug on the standard error of the mean (SEM) of prestimulus pupil size ($F_{2,54}$=8.13, p<.001), but no significant effect of task type ($F_{1,27}$=1.42, p=.24). Post-hoc t-tests indicated that the prestimulus pupil size distribution was significantly wider for ATX as compared to PLC (t(27)=3.43, p=.002; **Figure 1G**), as well as compared to DNP (t(27)=2.89, p=.007). DNP did not modulate the SEM of prestimulus pupil size significantly as compared to PLC (t(27)=1.97, p=.06). Taken together, these results suggest that ATX, but not DNP, increased arousal as indexed by prestimulus pupil size.

In addition to pupil size, we used heart rate (HR) and blood pressure (BP) to measure physiological arousal levels. For both these measures, we calculated the relative change of each measure as compared to a baseline measurement at the start of the session. We observed clear effects of the drugs on HR ($F_{2,54}$=8.36, p<.001) and BP ($F_{2,54}$=8.05, p<.001). Post-hoc tests indicated that ATX increased both HR and BP as compared to PLC (HR: t(27)=4.11, p<.001, **Figure 1H**; BP: t(27)=4.33, p<.001, **Figure 1I**), but DNP did not (HR: t(27)=1.01, p=.32; BP: t(27)=1.76, p=.09). All in all, our physiological measures are in line with the pupil data, suggesting that ATX increased arousal, whilst DNP did not.

**Supplementary Table 1**

*Mean prestimulus pupil size across tasks and for each task type separately*

| | All | Discrimination | Detection |
|---|---|---|---|
| PLC | 3.44 | 3.44 | 3.44 |
| DNP | 3.40 | 3.37 | 3.43 |
| ATX | 3.90 | 3.86 | 3.94 |

*Note.* Prestimulus pupil size is expressed in recorded pixels divided by 1000. As the number of recorded pixels depends on the settings of the eye tracker, the reported values can be considered as arbitrary units.

**Effects of the pharmacological manipulations and decision types on task performance**

Even though we were primarily interested in the influence of the pharmacological manipulations on the shape of the arousal-performance relationship, we also assessed the effects of the drugs and tasks on overall performance (d' and RT).

The pharmacological manipulations did not affect d' ($F_{1,54}$=1.05, p=.36), nor did the decision type ($F_{1,27}$=0.52, p=.48). The interaction effect between drugs and decision type on sensitivity was also not significant ($F_{1,54}$=2.79, p=.07). Sensitivity on the discrimination tasks was significantly higher after ingestion of ATX as compared to PLC (t(27)=2.42, p=.02), but there was no significant change in sensitivity after ingestion of DNP as compared to PLC (t(27)=1.34, p=.19). The drugs did not appear to affect sensitivity on the detection tasks (ATX vs. PLC: t(27)=0.06, p=.95; DNP vs. PLC: t(27)=0.33, p=.74).

The drugs did not appear to affect RTs ($F_{1,54}$=0.93, p=.40), but decision type did influence RTs significantly ($F_{1,27}$=17.51, p<.001), with smaller RTs for the detection tasks. The interaction effect between drug and decision type on RTs was not significant ($F_{1,54}$=0.72, p=.49). Within the discrimination tasks, there were no significant drug effects on RT (ATX vs. PLC: t(27)=0.83, p=.42; DNP vs. PLC: t(27)=0.49, p=.63). Within the detection tasks, there were also no significant effects of the drugs on RT (ATX vs. PLC: t(27)=1.13, p=0.27; DNP vs. PLC: t(27)=1.42, p=.17).

**Supplementary Table 2**

*The relationship between arousal and performance is quadratically shaped across pharmacological manipulations*

| | PLC | | DNP | | ATX | |
|---|---|---|---|---|---|---|
| | ∆AIC | ∆BIC | ∆AIC | ∆BIC | ∆AIC | ∆BIC |
| d' | 22.9 | 18.6 | 32.8 | 28.5 | 37.4 | 33.1 |
| RT | 20.1 | 15.8 | 30.1 | 25.8 | 36.6 | 32.2 |

*Note.* ∆AIC and ∆BIC are calculated by subtracting AIC (BIC) of the quadratic model from the AIC (BIC) of the linear model. ∆ values larger than 10 indicate that the quadratic model is favorable. These model fits are performed for the data pooled across tasks.

**The arousal-performance relationship is quadratically shaped for detection and discrimination decisions**

After having assessed that the relationship between arousal and performance was inverted U-shaped across decision types for all pharmacological agents (see **Supplementary Table 1**), we assessed the shape of the pupil-performance relationship for each decision type separately. To compensate for the lowered statistical power after splitting the data according to task type, we now used five pupil bins combined with Polynomial regression (see **Methods**). We test the coefficients of the linear ($ß_1$) and quadratic ($ß_2$) models against zero using one-sample t-tests. Because our aggregated data indicated that the arousal-performance relationship was quadratically shaped, we performed one-tailed tests for the quadratic models. As before, we used d' and RT as the dependent measures of performance. We first describe all results for d' (**Figure 2B-C**), followed by RT (**Figure 2E-F**).

For the discrimination tasks in the PLC condition, neither the quadratic or linear model appeared to explain the relationship between pupil size and sensitivity ($ß_2$: $t(27)=-1.35$, $p=.09$; $ß_1$: $t(27)=0.89$, $p=.38$; **Figure 2B**). D' was highest at intermediate levels of pupil-linked arousal for the detection tasks under PLC ($ß_2$: $t(27)=-2.18$, $p=.04$), although there was also evidence for a negative linear relationship ($ß_1$: $t(27)=-1.84$, $p=.04$; **Figure 2C**). For the DNP condition, in which arousal appeared to be unmodulated as compared to PLC, the pupil-performance relationship appeared to be quadratic for the discrimination tasks ($ß_2$: $t(27)=-4.75$, $p<.001$; **Figure 2B**), as well as for the detection tasks ($ß_2$: $t(27)=-2.19$, $p=.02$, but also $ß_1$: $t(27)=-2.43$, $p=.02$; **Figure 2C**). For the ATX condition, the pupil-performance relationship also appeared to be quadratic for both the discrimination tasks ($ß_2$: $t(27)=-4.19$, $p<.001$; **Figure 2B**) and the detection tasks ($ß_2$: $t(27)=-2.66$, $p=.007$; **Figure 2C**).

The RT results point towards U-shaped relationships between prestimulus pupil and RT for all drugs and decision types. For the PLC condition, RTs were shortest at intermediate pupil sizes for the discrimination tasks ($ß_2$: $t(27)=2.09$, $p=.02$; **Figure 2E**) as well as the detection tasks

($ß_2$: t(27)=3.13, p=.002; **Figure 2F**). Similarly, for the DNP condition, the pupil-RT relationship appeared to be U-shaped for the discrimination tasks ($ß_2$: t(27)=2.73, p=.006; **Figure 2E**) and the detection tasks ($ß_2$: t(27)=2.66, p=.007; **Figure 2F**). Lastly, the relationship between pupil-linked arousal and RT also appeared to be quadratically shaped under ATX during the discrimination tasks ($ß_2$: t(27)=2.50, p=.009; **Figure 2E**) and the detection tasks ($ß_2$: t(27)=2.11, p=.02; **Figure 2F**).

**Pharmacodynamics and pharmacokinetics of atomoxetine and donepezil**

The pharmaceuticals used in this study were chosen on the basis of their pharmacokinetic and pharmacodynamic properties, the relatively limited side-effects, and prior use of these pharmaceuticals in other studies in the cognitive sciences[1–4]. Atomoxetine is a relatively selective noradrenaline reuptake inhibitor, which inhibits the presynaptic noradrenaline reuptake transporter, thereby resulting in increased noradrenaline and dopamine levels in the synaptic cleft[5]. The half-life of atomoxetine varies between 4.5-19 hours and peak plasma levels are reached ~2 hours after administration. Donepezil is a cholinesterase inhibitor, which impedes breaking down of acetylcholine by cholinesterase thereby resulting in increased acetylcholine levels in the synaptic cleft. The elimination half-life of donepezil is 70 hours and peak plasma levels are reached after ~4 hours[6].

**Inclusion and exclusion criteria for participation in pharmacological study**

Participants were only allowed to participate if they met all the following criteria:

- Male
- 18-30 years old
- Right-handed
- Native Dutch-speaking
- BMI > 18.5 and <30
- Non-smoker
- No more than 15 alcoholic consumptions per week
- No (recreational) drug use (<1 time per month was allowed)
- No first line family (e.g. mother, brother, child) diagnosed with mental illness No known physical or mental illnesses (requiring regular use of medication)

**References**


1. Gratton, C., Yousef, S., Aarts, E., Wallace, D.L., D'Esposito, M., and Silver, M.A. (2017). Cholinergic, But Not Dopaminergic or Noradrenergic, Enhancement Sharpens Visual Spatial Perception in Humans. J. Neurosci. *37*, 4405–4415.

2. Boucart, M., Bubbico, G., Szaffarczyk, S., Defoort, S., Ponchel, A., Waucquier, N., Deplanque, D., Deguil, J., and Bordet, R. (2015). Donepezil increases contrast sensitivity for the detection of objects in scenes. Behav. Brain Res. *292*, 443–447.

3. Pfeffer, T., Ponce-Alvarez, A., Tsetsos, K., Meindertsma, T., Gahnström, C.J., van den Brink, R.L., Nolte, G., Engel, A.K., Deco, G., and Donner, T.H. (2021). Circuit mechanisms for the chemical modulation of cortex-wide network interactions and behavioral variability. Science Advances *7*, eabf5620.

4. Pfeffer, T., Avramiea, A.-E., Nolte, G., Engel, A.K., Linkenkaer-Hansen, K., and Donner, T.H. (2018). Catecholamines alter the intrinsic variability of cortical population activity and perception. PLoS Biol. *16*, e2003453.

5. Simpson, D., and Plosker, G.L. (2004). Atomoxetine: a review of its use in adults with attention deficit hyperactivity disorder. Drugs *64*, 205–222.

6. Rogers, S.L., and Friedhoff, L.T. (1998). Pharmacokinetic and pharmacodynamic profile of donepezil HCl following single oral doses. Br. J. Clin. Pharmacol. *46 Suppl 1*, 1–6.
